# Much Ado About Nothing: Accelerating Maximum Likelihood Phylogenetic Inference via Early Stopping to evade (Over-)optimization

**DOI:** 10.1101/2024.07.04.602058

**Authors:** Anastasis Togkousidis, Alexandros Stamatakis, Olivier Gascuel

**Affiliations:** Computational Molecular Evolution Group, Heidelberg Institute for Theoretical Studies, Germany; Institute of Theoretical Informatics, Karlsruhe Institute of Technology, Karlsruhe, Germany; Biodiversity Computing Group, Institute of Computer Science, Foundation for Research and Technology – Hellas; Institut de Systématique, Evolution, Biodiversité (ISYEB, UMR7205 - CNRS, Muséum National d’Histoire Naturelle, SU, EPHE, UA), Paris, France

**Keywords:** Early stopping, Stopping criteria, Maximum Likelihood

## Abstract

Maximum Likelihood (ML) based phylogenetic inference constitutes a challenging optimization problem. Given a set of aligned input sequences, phylogenetic inference tools strive to determine the tree topology, the branch-lengths, and the evolutionary parameters that maximize the phylogenetic likelihood function. However, there exist compelling reasons to not push optimization to its limits, by means of early, yet adequate stopping criteria. Since input sequences are typically subject to stochastic and systematic noise, one should exhibit caution regarding (over-)optimization and the inherent risk of overfitting the model to noisy input data. To this end, we propose, implement, and evaluate four statistical early stopping criteria in RAxML-NG that evade excessive and compute-intensive (over-)optimization. These generic criteria can seamlessly be integrated into other phylogenetic inference tools while not decreasing tree accuracy. The first two criteria quantify input data-specific sampling noise to derive a stopping threshold. The third, employs the Kishino-Hasegawa (KH) test to statistically assess the significance of differences between intermediate trees *before*, and *after* major optimization steps in RAxML-NG. The optimization terminates early when improvements are insignificant. The fourth method utilizes multiple testing correction in the KH test. We show that all early stopping criteria infer trees that are statistically equivalent compared to inferences without early stopping. In conjunction with a necessary simplification of the standard RAxML-NG tree search heuristic, the average inference times on empirical and simulated datasets are ∼3.5 and ∼1.8 times faster, respectively, than for standard RAxML-NG v.1.2. The four stopping criteria have been implemented in RAxML-NG and are available as open source code under GNU GPL at https://github.com/togkousa/raxml-ng.

## 1. Introduction

Phylogenetic Inference addresses the problem of finding a binary tree which optimally, according to some optimality criterion, represents the evolutionary relationships among biological sequences, provided in the form of a Multiple Sequence Alignment (MSA). There exist a plethora of optimality criteria to infer trees, such as Maximum Parsimony (MP) (Fitch, 1971), Maximum Likelihood (ML) (Felsenstein, 1981), or the least squares method (Cavalli-Sforza and Edwards, 1967; Fitch and Margoliash, 1967). ML tree inference constitutes a computationally challenging optimization problem. It strives to find *the* optimal tree topology (a discrete parameter), along with the corresponding optimal branch lengths and evolutionary model parameters (continuous parameters) that maximize the phylogenetic likelihood score. The likelihood reflects the probability that the observed sequences have evolved under the specific binary tree topology and the respective continuous model parameters (Yang, 2014). Software tools such as RAxML (Stamatakis, 2014), RAxML-NG (Kozlov et al., 2019), IQ-TREE (Minh et al., 2020), and PhyML (Guindon et al., 2010) deploy thorough, hill-climbing heuristics (St. John, 2016) in their attempt to find the tree and parameters that maximize the likelihood function. The results of these tools are typically local optima as the tree inference problem under ML has been shown to be NP-hard (Roch, 2006).

Sequences, however, are subject to noise (Townsend et al., 2012) stemming from both, stochastic, and systematic sources. Evolution, which is stochastic in nature, induces a form of *stochastic noise* in the sequences, onto which *Sampling Noise* is superimposed (Hillis and Huelsenbeck, 1992; Münkemüller et al., 2012). Stochastic noise reflects the extent to which the observed data distribution deviates from the theoretically expected values. Sampling Noise occurs because the sequences typically used for a phylogenetic analysis merely represent a small fraction of the corresponding genomes. Thus, the appropriateness of such a sample to approximate the original distribution may be put into question. Further, systematic noise, that is predominantly prevalent in empirical MSAs, may be caused by a plethora of phenomena such as sequencing errors (Moutsopoulos et al., 2021), alignment errors (Dress et al., 2008), homoplasy (Rokas and Carroll, 2006), and gene tree - species tree discordance in supermatrix MSAs (Maddison, 1997).

Given this intrinsic noise in sequence data, exhaustive optimization might lead to overfitting, wherein the inferred statistical model fits both, the phylogenetic signal *and* noise in the MSA. A systematic study which examines overfitting of ML inference tools on both, empirical, and simulated MSAs (de Vienne et al., 2017) concludes that a substantial part of the optimization steps in RAxML and PhyML is unnecessary and can be ommited via integration of appropriate early stopping criteria for ML inference. For example, a straight-forward approach is to employ more superficial, yet significantly faster, tree search heuristics, such as implemented in FastTree (Price et al., 2010). However, a recent benchmark study (Höhler et al., 2022) indicates that FastTree is outperformed by more thorough heuristics (IQ-Tree, RaxML-NG) with respect to the statistical plausibility of the inferred trees. Another study focusing on the reconstruction of the tree of mammals (Lemoine et al., 2018) shows that RAxML-NG infers a phylogeny with substantially higher bootstrap support values (Felsenstein, 1985) than FastTree. Therefore, the objective of our study is to integrate and apply reasonable ML inference stopping criteria to circumvent unnecessary optimization steps, without compromising the quality of the inferred tree.

To ascertain convergence and terminate the execution, RAxML-NG, PhyML, and FastTree use small, yet arbitrarily chosen log-likelihood improvement thresholds. For instance, RAxML and RAxML-NG v.1.1 use a default improvement threshold of 0.1 log-likelihood units. Based on the results of a systematic study that quantifies the impact of varying numerical thresholds on ML tree inference (Haag et al., 2023), the default threshold value in RAxML-NG v.1.2 was increased to 10 log-likelihood units. In the current study we argue that these numerical thresholds should not be universally fixed, but rather dynamically adjusted to the specific phylogenetic signal of the dataset at hand, the inherent noise in the MSA, and, potentially, to the convergence dynamics of each independent tree inference. While users can modify the convergence threshold values in standard, they often lack the patience, time, and knowledge to select the most appropriate threshold for their dataset. This emphasizes the need for automated dataset-specific stopping criteria, such as those we propose here.

Along these lines, IQPNNI (Vinh and Von Haeseler, 2004), the predecessor of IQ-TREE, employs a more sophisticated approach to terminate the tree search. This stopping rule statistically evaluates the time intervals (in terms of number of iterations) between successive occurrences of tree topologies which improve the log-likelihood score; the tree search terminates when the estimated probability (using a Weibull distribution of record values) that the optimal tree topology has been found exceeds 95%. However, this approach was abandoned in later versions of IQ-TREE. The current version of IQ-TREE terminates when no better tree is found after a predefined number of iterations (default value *N* := 100).

In our study we propose four early stopping criteria that dynamically adjust the log-likelihood improvement threshold (*ϵ*) value by taking into account the inherent noise in the data as well as the convergence dynamics of each individual Maximum Likelihood (ML) tree inference. We implement these stopping criteria in RAxML-NG v.1.2. Importantly, these criteria are sufficiently generic such that they can seamlessly be integrated into other phylogenetic inference tools that deploy numerical thresholds for convergence assessment such as PhyML, FastTree, and IQ-TREE. Our experimental results on 725 representative empirical datasets (575 DNA and 150 amino-acid (AA) MSAs), and representative 506 simulated (DNA) MSAs, show that, when tree inferences are initiated on a parsimony starting tree early stopping yields trees which are statistically equivalent to those inferred under the default constant ad hoc convergence threshold in 99% of the cases. Furthermore, in conjunction with a necessary simplification of the standard RAxML-NG search strategy (see Section 2), the average inference times for empirical and simulated datasets are ∼3.5 and ∼1.8 times faster, respectively, than those of standard RAxML-NG v.1.2. All MSAs we used for our experiments, are available for download at https://cme.h-its.org/exelixis/material/stopping_criteria.tar.gz.

## 2. Materials and Methods

In this section, we initially introduce an alternative, simplified version of RAxML-NG. We use this simplified version to develop and assess our methods. Next, we present four distinct, fast-to-compute, stopping criteria for early stopping in ML tree inference based on the rationales discussed in the preceding section. The first two criteria strive to quantify the inherent Sampling Noise (SN) in the input MSA. Specifically, the first method assumes a Normal Distribution to approximate Sampling Noise (SN-Normal), while the second employs a non-parametric RELL-like (Kishino et al., 1990) approach (SN-RELL). The third criterion relies on the Kishino-Hasegawa (KH) test (Kishino and Hasegawa, 1989) to statistically assess significant log-likelihood differences between tree topologies *before* and *after* Subtree Prune and Regraft (SPR) Rounds, which are the core topological optimization steps in RAxML-NG. We terminate early when improvements between such subsequent tree topologies are statistically insignificant. The fourth criterion incorporates a multiple testing correction for the KH test.

### 2.1 Simplified RAxML-NG (sRAxML-NG)

When the improvement in log-likelihood drops below a predefined, fixed *ϵ*-threshold in RAxML-NG, the ML tree inference does not terminate immediately, so as to potentially navigate out of local optima. Instead of terminating, RAxML-NG continues the tree search by adjusting SPR-specifc parameters for the subsequent SPR rounds (e.g., it increases the maximum SPR radius or switches from superficial (FAST) to more thorough (SLOW) SPR rounds), after conducting a round of Model Parameter Optimizations (MPO). The main difference between FAST and SLOW SPR rounds is that, during FAST SPR rounds, RAxML-NG evaluates each tree topology generated by an SPR move using the existing branch lengths, while in SLOW SPR rounds, the lengths of the three adjacent branches around the insertion node are re-optimized. Further, full Branch-Length Optimization (BLO) rounds are conducted after each FAST or SLOW SPR round and prior to MPO. Hence, the tree search only terminates after several unsuccessful iterations of SLOW SPR rounds with distinct SPR parametrizations (Kozlov, 2018).

Within this complex optimization framework, the impact of convergence thresholds is challenging to assess, as termination does not occur immediately when the score improvement drops below the predefined, fixed *ϵ* value. As a consequence, it is difficult to evaluate and compare the impact of distinct stopping criteria that directly modify the *ϵ* value in an objective manner. Therefore we initially simplified and accelerated, this complex default search strategy in RAxML-NG which constitutes a first form of early stopping. We call this alternative search heuristic *Simplified RAxML-NG (sRAxML-NG)*. sRAxML-NG conducts a sequence of FAST SPR Rounds followed by a sequence of SLOW SPR rounds. Each sequence terminates when the log-likelihood improvement drops below a static (as in standard RAxML-NG) or dynamic (as defined by our criteria) *ϵ* threshold. In contrast to the standard RAxML-NG search strategy, we simply execute the FAST and SLOW SPR rounds with a fixed maximum subtree re-insertion radius parameter of 10. Full BLO rounds are conducted after each FAST and each SLOW SPR round, as in standard RAxML-NG. MPO rounds are conducted on the initial tree topology (after BL0), in-between the sequences of FAST and SLOW SPR rounds, and on the final tree (before termination). This simplified heuristic substantially speeds up inferences. This speedup is a positive by-product of our study as sRAxML-NG was predominantly developed to facilitate studying the impact and dynamic adaptation of the *ϵ* convergence thresholds. Further details and workflow diagrams for the standard RAxML-NG and the sRAxML-NG heuristics are provided in the Supplementary Material.

### 2.2 Sampling Noise Normal Distribution (SN-Normal)

In the case of sampling noise, the “true” log-likelihood value is the expected log-likelihood value when drawing a sample of size *s* (i.e., the number of sites in the MSA) from a from a substantially larger (full genome) MSA. The observed variations around the true value correspond to sampling noise and allow to quantify the ability of the sample to approximate the complete distribution. We adopt a simple model wherein we posit that the log-likelihood is subject to a type of noise that is invariant *before* and *after* the fundamental optimization block (SPR round) and that it follows the same normal distribution (SN-Normal). In other words:

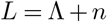

and:

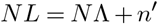

Here, *n* and *n*^*′*^ denote independent noise drawn from the same distribution, Λ and *N* Λ denote the “true” log-likelihood values corresponding to the trees before and after an SPR round, respectively, and *L* and *NL* denote the noisy log-likelihood values for the same tree topology (*NL* ≥ *L*). By assuming independence between *n* and *n*^*′*^, we can estimate the parameters of this distribution (i.e., the mean and standard deviation, assuming normality) merely using a single, yet reasonable, that is non-random (e.g., parsimony) starting tree. We propose the following straight-forward parametric solution:

1. Consider the set of per-site log-likelihood values for the *s* sites in the MSA. Their sum is *L*, and the standard deviation is *σ*_*SN*_.
2. The standard deviation of the sum over the per-site log-likelihoods (equal to *L*) is 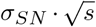.
3. Under these parametric assumptions, the quan-tity 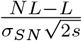 follows a normal distribution *N* (0, 1). We can derive the 95% confidence interval via the cumulative distribution function Φ(•) of a standard normal distribution. We continue the search with 95% confidence, if:

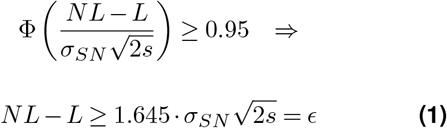

This *ϵ* value is based on a normal assumption and uses the central limit theorem because we sum over a sufficiently large number of independent variables (i.e., persite log-likelihoods). The simplicity of this approach allows for rapid computation of the *ϵ*-threshold, while maintaining reasonable accuracy, yet under certain conditions only, as we show via our experiments.

### 2.3 Sampling Noise RELL Approximation (SN-RELL)

In analogy to the SN-Normal stopping rule, we also propose a non-parametric version via the RELL bootstrap approach (SN-RELL), as follows:

1. Resample *s* per-site log-likelihoods *with* replacement from a reasonable (e.g., parsimony) starting tree, and compute the sum 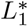 over these *s* values.
2. Repeat the experiment in step 1 *N* times (e.g., *N* := 1, 000) to obtain *N* 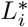, *i* = 1, 2, …, *N* values. The non-parametric, empirical distribution of 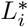 values represents the variability of *L* and *NL*. In the next step we estimate the distribution of *NL* − *L* assuming independence of *NL* and *L* and the Null Hypothesis being *N* Λ = Λ.
3. Randomly and independently draw *M* (e.g., *M* =: 1, 000) pairs of 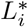 values, where *NL*^*^ and *L*^*^ denote the maximum and minimum of a single pair, and represent *NL* and *L*, respectively. Compute *NL*^*^ − *L*^*^ and the 95% quantile of the distribution of *M* differences of random pairs. This yields an *ϵ* threshold analogous to the one we propose in (eq. 1), but is non-parametric. In other words, when *NL* − *L* exceeds the 95% quantile of the distribution of differences, we are confident that the optimization step significantly improved the log-likelihood score. Otherwise, the difference *NL* − *L*, that is the log-likelihood difference cannot be distinguished from Sampling Noise.

### 2.4 Kishino-Hasegawa (KH) test

The independence assumptions between *NL* and *L* underlying the proposed Sampling Noise methods (Sections 2.2 and 2.3) might be strong and not necessarily valid. The *ϵ* values generated by the two methods are comparatively large in practice, of the order of thousands of log-likelihood units (see Supplement, Figure S8). Based on our experience with developing ML heuristics and the results of Haag *et al*. (2023), these *ϵ* values might result in stopping too early and, hence, caution is warranted when employing such thresholds. We elaborate further on these limitations in Sections 3 and 4, as well as in the Supplementary Material.

Thus, we also deploy the KH test to compare subsequent best trees before and after each SPR round. Based on the per-site log-likelihood differences of the two trees, an *ϵ* value can be derived, which depends on the variations observed in the per-site log-likelihood distributions corresponding to the SPR round under consideration. Consequently, the *ϵ* threshold will vary across SPR rounds and automatically adapts to the convergence dynamics of each independent ML tree search. The main difference with respect to the SN-based approaches outlined above, is that we do account for the dependence between *n* and *n*^*′*^ (noise before and after an SPR round), and consequently between *L* and *NL*. With the KH test, we use a form of paired test, while with the SN-based approaches, the tests are unpaired. More specifically, for an MSA with *s* sites, suppose that the per-site log-likelihood vectors before (*V*_*L*_) and after (*V*_*NL*_) any given SPR round, are the following:

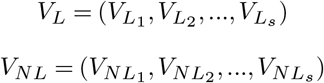

Where:

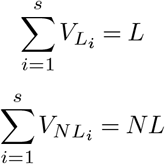

The KH test can be applied as follows:

1. Compute the distribution of the (paired) differences between the vector coordinates 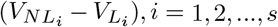. These differences can be either positive or negative, even if *NL−L >* 0.
2. Compute the standard deviation *σ*_*KH*_ of the distribution of the *s* differences. The standard deviation of the difference *NL−L* is equal to 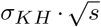.
3. We assume that the quantity 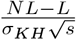 follows a normal distribution *N* (0, 1). We continue the tree search with 95% confidence if:

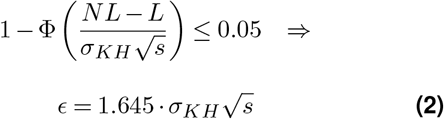

### 2.5 Kishino-Hasegawa (KH) test with multiple correction

We deem the assumptions underlying the KH test to be more acceptable and realistic than those of the SN-based approaches. While the latter might induce premature termination due to their relatively large thresholds, the KH test may unnecessarily prolong the tree search. This is primarily because the test does not incorporate adjustments for multiple testing. In fact, in an SPR round we evaluate the likelihood of many potential SPR topologies (i.e., topologies generated from multiple SPR moves which constitute a SPR round) and only select the best one. This is typically a situation that requires accounting for the multiplicity of the tests. However, the majority of generated topologies during an SPR round merely decrease the fit of the tree (the likelhood) to the input MSA, especially toward the end of the optimization process when we are close to the (local) optimum. To correct for multiple testing, we count the number *N*_*t*_ of SPR topologies which improve the likelihood of the best tree prior to initiating the SPR round, and apply a Bonferroni correction to the p-value of the output tree, that is, the tree after termination of the SPR round. The adjusted *ϵ* threshold is then derived as follows:

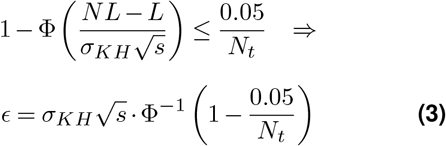

## 3. Results

We conducted experiments on 725 empirical and 506 simulated MSAs. Among the empirical datasets, 575 are DNA datasets and 150 are AA MSAs, whereas all simulated MSAs are DNA datasets, in order to reduce the CO_2_ footprint. We sampled the empirical MSAs from the TreeBASE databse (Piel et al., 2009) and the simulated MSAs from the datasets used in a recent benchmark study conducted by our group and colleagues (Trost et al., 2024). The selected empirical and simulated MSAs capture the full spectrum of difficulty scores predicted by the Pythia tool (Haag et al., 2022). Given the similarity in the results between empirical DNA and AA datasets, we only present the findings from empirical and simulated DNA MSAs for the sake of readability. Detailed results for AA MSAs, along with additional information on dataset sizes, Pythia score distributions, and the evolutionary models used in the analysis, can be found in the Supplementary Material.

We refer to all RAxML-NG versions proposed in Section 2, including the sRAxML-NG version, as the *Early Stopping* versions/criteria. In our experimental setup, we compare the output of RAxML-NG v.1.2 with the Early stopping versions. For each MSA, we conduct two experiments per RAxML-NG version, one using the same set of 10 distinct parsimony starting trees for all versions (i.e., trees with a “good” parsimony score) and a second one using the same set of 10 distinct random starting trees for all versions. Therefore, each version conducts 10 independent tree inferences in every execution. We refer to the 10 output trees from each execution as the inferred *ML trees*, while we denote the tree with the highest likelihood score among all 10 ML trees per RAxML-NG version as the *best ML tree*. The best ML tree inferred via RAxML-NG v.1.2 will be the *best standard RAxML-NG tree*.

Our benchmark comparisons include (*i*) plausibility assessment on the ML trees inferred via the early stopping versions, (*ii*) topological accuracy assessment of the best ML trees inferred by each version against some reference tree (see below), and (*iii*) execution time measurements. To assess plausibility, we conduct pairwise comparisons between all ML trees inferred from the Early Stopping versions with the best standard RAxML-NG tree using the KH test. If Null Hypothesis is accepted, we conclude that the two trees are statistically equivalent, and therefore the ML tree inferred via early stopping is *plausible*. If the Null Hypothesis is rejected, and the ML tree inferred via early stopping has a lower log-likelihood score, we consider it to be statistically worse (implausible) compared to the best standard RAxML-NG tree. There are rare instances, though, where the Null Hypothesis is rejected, yet the log-likelihood score of the ML tree inferred via early stopping is higher; in such cases we consider the early stopping methods to have yielded a statistically better ML tree. Regarding topological accuracy, we compare the best ML trees inferred from all versions with the reference tree using the Robinson–Foulds (RF) distance (Robinson and Foulds, 1981). In simulated data, the reference tree is the “true” tree used to generate the data, while on empirical MSAs we use the best standard RAxML-NG tree. Finally, to attain accurate runtime measurements, we measure the speedups gained of early stopping by executing *all* versions sequentially. Note that an efficient parallelization of our stopping criteria is already implemented and tested in the respective open source code.

We ran our experiments on the Haswell/URZ Cluster, located at the Computing Center of the University of Heidelberg. It consists of 224 nodes with Intel Haswell CPUs (E5-2630v3 running at 2.40GHz). Each node contains 2 CPUs and each CPU has 8 physical cores. The operating system is CentOS Linux 7 (Core).

Figure 1 illustrates the distribution of the number of statistically equivalent and worse ML trees inferred from Early Stopping versions on empirical DNA datasets. Figure 1a corresponds to executions using 10 parsimony starting trees, while 1b to inferences with 10 random starting trees. The rare instances where Early Stopping versions infer statistically better ML trees are reported below.

**Figure 1.**
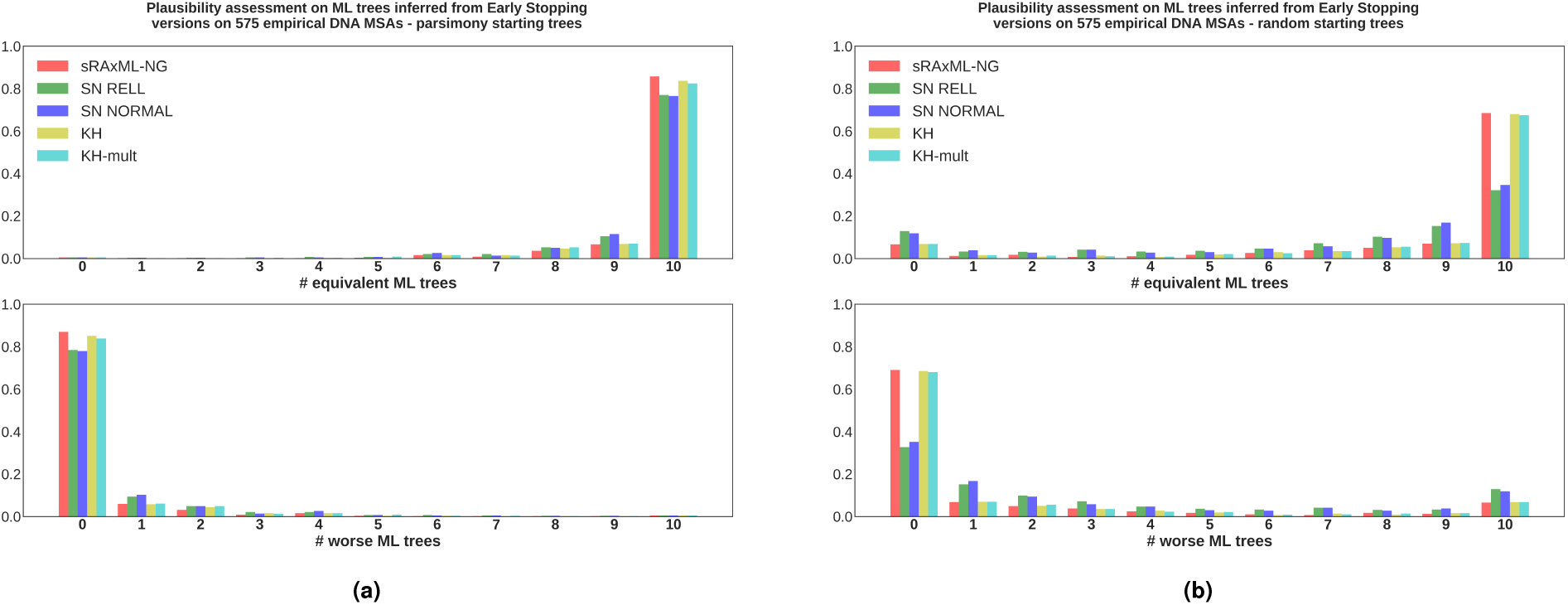
**(a)** Results from the plausibility tests conducted on the ML trees inferred from Early stopping criteria on 575 empirical DNA MSAs. For each dataset, all RAxML-NG versions conduct 10 independent tree inferences using (the same) set of parsimony starting trees. The ML trees inferred using Early Stopping criteria are compared to the best ML tree inferred from RAxML-NG v.1.2 using the KH test in a pair-wise manner. The horizontal axes shows the number (out of 10) of the ML trees inferred by each version that are statistically equivalent (top) and worse (bottom) than the best ML tree inferred via RAxML-NG v.1.2. The vertical axes shows the fraction of MSAs whose analysis yielded the corresponding number of equivalent or worse ML trees, respectively. **(b)** Identical experimental setup as under (a), with the sole difference being that we use 10 random starting trees instead of 10 parsimony starting tree.

Regarding parsimony starting trees (Figure 1a), all Early Stopping versions infer at least one plausible tree in 572 MSAs (99.4% of the cases). The SN-based methods exhibit slightly lower accuracy in terms of statistical plausibility compared to sRAxML-NG and the KH-based methods. Specifically, the number of MSAs in which the SN-based methods infer 10 plausible trees is 440 for the SN Normal (76%) and 443 for SN RELL version (77%), respectively. The corresponding numbers for the remaining methods are 493 for sRAxML-NG (86%), 481 for the simple KH test (84%), and 474 for the KH-multiple testing version (82%). The reduced accuracy of the SN-based methods is attributed to the large *ϵ* values they yield, even when computed on parsimony starting trees. These *ϵ*-thresholds, ranging from hundreds to thousands log-likelihood units (see Supplement, Figure S8), sometimes induce premature termination of the inference process. This limitation is more prevalent when using random starting trees (see Figure 1b). Further, regarding the rare instances where Early Stopping versions infer statistically better ML trees than the best standard RAxML-NG tree, the KH-multiple testing version infers at least one (out of the 10 ML trees) statistically better tree on 10 MSAs, while the remaining Early Stopping versions on 8 MSAs. This occurs because tree searches use ad hoc heuristics, and the search paths in the vast tree search space diverge at different stages of the search for each version, depending on the *ϵ* values in conjunction with the complexity of each search strategy. Consequently, although RAxML-NG v.1.2 employs a thorough heuristic, there is always a small probability that a superficial heuristic might follow a distinct search path, yielding an even statistically better ML tree and not merely a tree with a slightly better likelihood.

Regarding random starting trees (Figure 1b), in all Early Stopping versions, the distributions of plausible ML trees, and therefore the distributions of implausible (statistically worse) ML trees, spread more broadly across the range [0, 10]. This is again expected, given that the starting points of the tree inferences are sub-optimal (in fact, the worst possible), and the stopping criteria may occasionally terminate the tree searches prematurely. Additionally, the SN-based methods perform significantly worse than sRAxML-NG and the KH-based methods. As already reported, the *ϵ*-thresholds computed by the SN-based versions on random starting trees are substantially larger than those computed on parsimony starting trees (see Supplement, Figure S8). This outcome aligns with one of the basic assumptions for quantifying Sampling Noise, that is, that the starting tree should be “reasonable” with respect to the data (see Section 2.2). Consequently, SN-based methods should not use random starting trees.

More specifically, SN Normal version infers 10 plausible ML trees in 199 MSAs (35%) and the SN RELL in 185 MSAs (32%). These numbers are considerably higher for sRAxML-NG at 394 (69%), for the simple KH at 391 (68%) and for the KH-multiple testing version at 388 (67%). Further, in 93% of the cases, sRAxML-NG and the KH-based versions infer at least one plausible tree, while the corresponding fraction for SN-based methods is 88%. Finally, sRAxML-NG and the KH-based versions infer at least one statistically better tree (compared to RAxML-NG v.1.2) on 4 MSAs, and the SN-based versions on 3 MSAs. The analogous plot to Figure 1 for simulated DNA MSAs is presented in the Supplementary Material.

Figure 2 illustrates the distributions of RF distances between the best ML trees inferred from each RAxML-NG version and the reference tree topologies, when 10 parsimony starting trees are used. For empirical MSAs, it is challenging to draw conclusions about the distributions, since the true tree topology is unknown. Consequently, we compare the topologies inferred by the Early Stopping versions against the best topology inferred by RAxML-NG v1.2. As shown in the Figure, the results from this comparison indicate that, in the majority of cases, the relative RF distance between the two topologies is below 20%. The plot also demonstrates that for simulated data, there is no significant difference in the distributions of relative RF distances between RAxML-NG v.1.2 and the Early Stopping versions, when the best ML trees are compared to the true tree topology. In fact, all RF distance distributions exhibit the same average value of 0.13. Moreover, this average value remains unaltered when random starting trees are used (see Supplement, Figure S7). We conclude that, in terms of topological accuracy, the Early Stopping versions do not perform worse than the standard RAxML-NG version.

**Figure 2.**
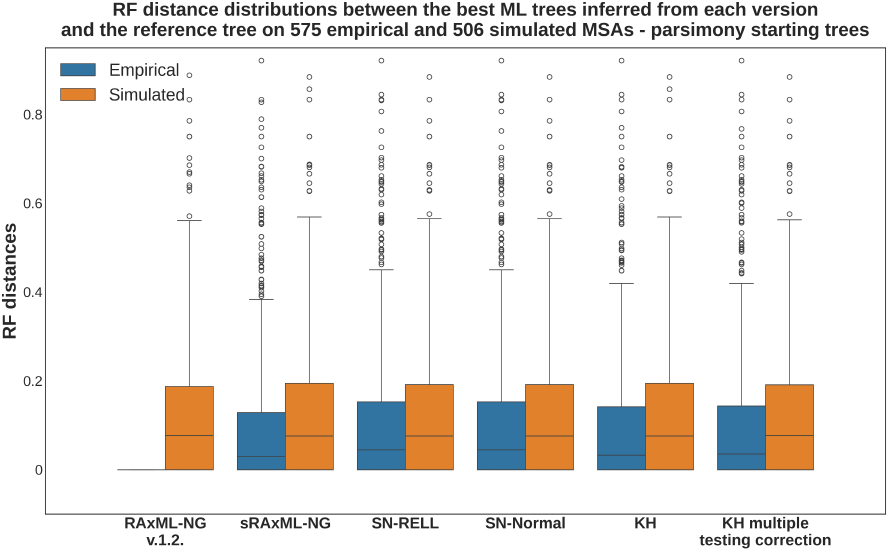
Distributions of relative RF distances between the best ML trees inferred from each RAxML-NG version and the corresponding reference tree topologies, where each version conducted 10 independent tree inferences on 575 empirical and 506 simulated DNA MSAs using parsimony starting trees. On simulated MSAs, the reference tree topology is the “true” tree used to generate the given sequences. For empirical MSAs, the reference topology is the best ML tree inferred by RAxML-NG v.1.2, explaining the absence of a distribution for empirical MSAs in the RAxML-NG v.1.2 column.

Finally, Figure 3 provides the runtime improvements of the Early Stopping versions relative to RAxML-NG v1.2, when 10 parsimony starting trees are used. Notably, the speedups observed on empirical MSAs are substantially higher than those on simulated MSAs. This outcome was expected, as simulated data are generally easier to analyze and tend to converge faster as they exhibit clearer and stronger signal. For empirical MSAs, the average speedup ranges from 3x for sRAxML-NG to 4x for the SN-based versions. The average speedups for the KH-based versions are 3.4x for the simple KH test and 3.6x for the KH test with multiple testing correction. Clearly, the highest contribution to the runtime improvement is due to the simplified RAxML-NG heuristic (sRAxML-NG). However, as indicated in Figure 4, stopping criteria further increase the speedup while maintaining tree inference accuracy. In the latter Figure we present the speedups of the SN-based and KH-based versions compared to the sRAxML-NG version. SN-based methods are approximately 30% faster, while KH-based methods are 13% (simple KH) and 20% (KH-multiple testing) faster, than sRAxML-NG.

**Figure 3.**
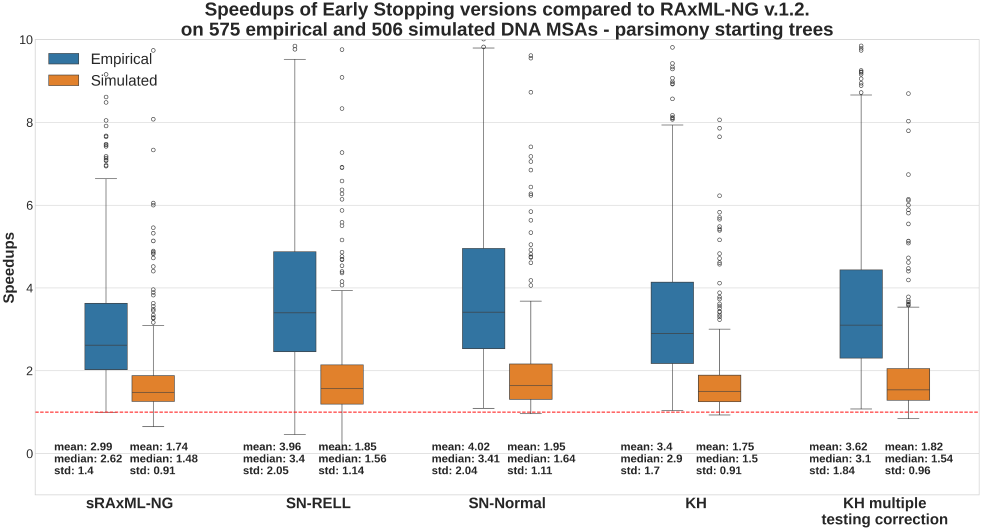
Speedup distributions of Early Stopping versions relative to RAxML v.1.2 on 575 empirical and 506 simulated DNA MSAs. The speedups refer to runtimes measured on sequential executions using parsimony starting trees. The dashed line at the bottom corresponds to a speedup of 1x.

**Figure 4.**
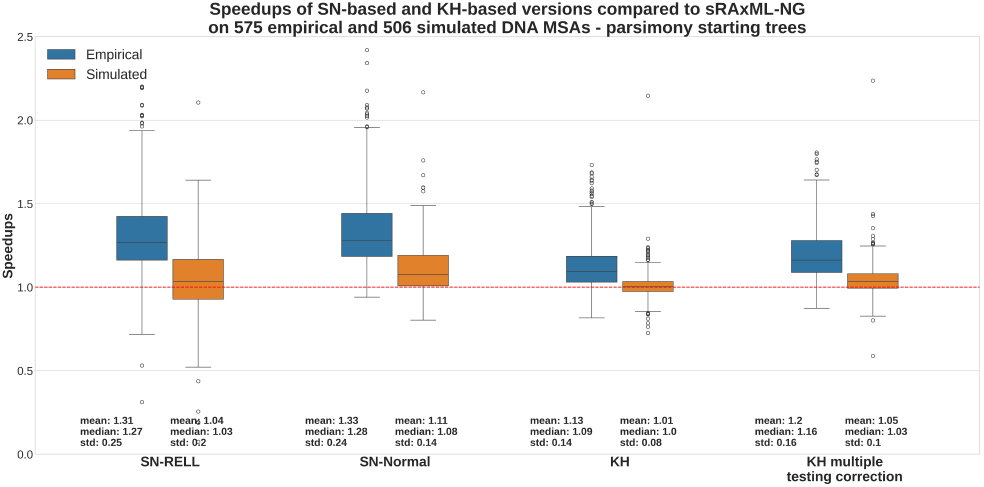
Speedup distributions of SN-based and KH-based RAxML-NG versions relative to sRAxML-NG on 575 empirical and 506 simulated DNA MSAs. The speedups refer to runtimes measured on sequential executions using parsimony starting trees. The dashed line at the bottom corresponds to a speedup of 1x.

The speedup and topological accuracy plots for searches initiated on random starting trees are analogous to Figures 2, 3, and 4. Corresponding plots are included in the Supplementary Material.

## 4. Discussion

We designed, implemented, and tested four statistical criteria for early termination in phylogenetic tree searches. These four methods are integrated into a new version of RAxML-NG called sRAxML-NG. This new version simplifies the tree search procedure and also performs adaptive early stopping in contrast to the standard RAxML-NG version, whose search strategy appears to be unnecessarily complex in retrospect and according to our experiments. The SN-based methods yield excessively large *ϵ*-values, inducing the highest average speedups at the price of reduced accuracy, especially when using random starting trees. Conversely, the KH-based methods are fast-to-compute and adaptive as they dynamically adjust the *ϵ*-threshold during each individual tree search. The multiple testing cor-rection in the KH test further refines this approach by accounting for the number of improving SPR topologies encountered during an SPR round.

Our comprehensive evaluation demonstrates that the proposed stopping criteria can be seamlessly integrated into phylogenetic inference heuristics, preserve tree inference quality, and yield runtime improvements between 3x and 4x. Notably, the KH-multiple testing criterion constitutes the most effective method as it yields an optimal balance between speed and accuracy. To better illustrate the impact of these improvements for practitioners, we present the absolute and relative run-time improvements achieved by the KH-multiple testing version across five large empirical MSAs in Table 1. We observe absolute runtime gains between 4 and 5 hours compared to the standard version and up to 50 minutes compared to sRAxML-NG without early stopping. Finally, in terms of topological accuracy, our Early Stopping versions performed are on par with RAxML-NG v.1.2 when assessed on simulated data.

**Table 1.** Table highlighting runtime improvements of Early Stopping using the KH-test with multiple testing correction on five large empirical MSAs. We compare both against RAxML-NG v.1.2 and the sRAxML-NG. The execution times of the three versions, shown in the third, fourth, and fifth column, correspond to 10 ML tree inferences using parsimony starting trees. The right-most column indicates the number of plausible trees inferred from Early Stopping using the KH-multiple testing version, as discussed in Figure 1.

| Dataset | Taxa, Sites | Standard time | sRAxML-NG time | KH-mult time | Speedup vs Standard | Speedup vs sRAxML-NG | Number of pl. trees |
| --- | --- | --- | --- | --- | --- | --- | --- |
| 26639_0 | 197, 72082 | 6h 22m | 1h 44m | 1h 14m | 5.1x | 1.4x | 10 |
| 26377_0 | 882, 3026 | 5h 5m | 43m | 29m | 10.4x | 1.5x | 5 |
| 1552_1 | 1205, 2114 | 4h 48m | 36m | 26m | 10.7x | 1.3x | 10 |
| 16630_0 | 1785, 807 | 4h 25m | 29m | 21m | 12.6x | 1.4x | 10 |
| 11872_1 | 169, 35603 | 5h 26m | 2h 20m | 1h 29m | 3.6x | 1.6x | 10 |

With respect to future work, we intend to leverage our understanding of the inherent complexity in analyzing a given MSA, as predicted by the Pythia score (Haag et al., 2022), along with our initial work on dynamically adapting the search strategy based on the Pyhtia score (Togkousidis et al., 2023) in conjunction with the stopping criteria we propose here. Our long-term objective will be to develop a new, fully adaptable version of RAxML-NG, which autonomously determines the number of starting trees, the thoroughness of the heuristic, the convergence dynamics, and the optimal point to terminate the search, based on the phylogenetic signal and behavior of the specific MSA to be analyzed.

## Acknowledgment

This study was financially supported by the Klaus Tschira Foundation and by the European Union (EU) under Grant Agreement No 101087081 (Comp-Biodiv-GR). O.G. was supported by PRAIRIE (ANR-19-P3IA-0001).

## Bibliography

Cavalli-Sforza, L. L. and Edwards, A. W. (1967). Phylogenetic analysis. models and estimation procedures. American journal of human genetics, 19(3 Pt 1):233.

de Vienne, D. M., Supek, F., and Gabaldon, T. (2017). Overtraining often results in topologically incorrect species trees with maximum likelihood methods. bioRxiv. Doi 10.1101/140780.

Dress, A. W., Flamm, C., Fritzsch, G., Grünewald, S., Kruspe, M., Prohaska, S. J., and Stadler, P. F. (2008). Noisy: identification of problematic columns in multiple sequence alignments. Algorithms for Molecular Biology, 3:1–10.

Felsenstein, J. (1981). Evolutionary trees from dna sequences: A maximum likelihood approach. Journal of Molecular Evolution, 17:368–376. doi: 10.1007/BF01734359.

Felsenstein, J. (1985). Confidence limits on phylogenies: an approach using the bootstrap. evolution, 39(4):783–791.

Fitch, W. M. (1971). Toward Defining the Course of Evolution: Minimum Change for a Specific Tree Topology. Systematic Biology, 20(4):406–416. doi: 10.1093/sysbio/20.4.406.

Fitch, W. M. and Margoliash, E. (1967). Construction of phylogenetic trees: a method based on mutation distances as estimated from cytochrome c sequences is of general applicability. Science, 155(3760):279–284.

Guindon, S., Dufayard, J.-F., Lefort, V., Anisimova, M., Hordijk, W., and Gascuel, O. (2010). New algorithms and methods to estimate maximum-likelihood phylogenies: assessing the performance of phyml 3.0. Systematic biology, 59(3):307–321.

Haag, J., Höhler, D., Bettisworth, B., and Stamatakis, A. (2022). From easy to hopeless—predicting the difficulty of phylogenetic analyses. Molecular Biology and Evolution, 39(12):msac254.

Haag, J., Hübner, L., Kozlov, A. M., and Stamatakis, A. (2023). The free lunch is not over yet—systematic exploration of numerical thresholds in maximum likelihood phylogenetic inference. Bioinformatics Advances, 3(1):vbad124.

Hillis, D. M. and Huelsenbeck, J. P. (1992). Signal, Noise, and Reliability in Molecular Phylogenetic Analyses. Journal of Heredity, 83(3):189–195. doi: 10.1093/oxfordjournals.jhered.a111190.

Höhler, D., Haag, J., Kozlov, A. M., and Stamatakis, A. (2022). A representative performance assessment of maximum likelihood based phylogenetic inference tools. bioRxiv. doi: 10.1101/2022.10.31.514545.

Kishino, H. and Hasegawa, M. (1989). Evaluation of the maximum likelihood estimate of the evolutionary tree topologies from dna sequence data, and the branching order in hominoidea. Journal of molecular evolution, 29:170–179.

Kishino, H., Miyata, T., and Hasegawa, M. (1990). Maximum likelihood inference of protein phylogeny and the origin of chloroplasts. Journal of Molecular Evolution, 31:151–160.

Kozlov, A. Models, Optimizations, and Tools for Large-Scale Phylogenetic Inference, Handling Sequence Uncertainty, and Taxonomic Validation. PhD thesis, Karlsruhe Institute of Technology, (2018).

Kozlov, A. M., Darriba, D., Flouri, T., Morel, B., and Stamatakis, A. (2019). RAxML-NG: a fast, scalable and user-friendly tool for maximum likelihood phylogenetic inference. Bioinformatics, 35(21):4453–4455. doi: 10.1093/bioinformatics/btz305.

Lemoine, F., Domelevo Entfellner, J.-B., Wilkinson, E., Correia, D., Dávila Felipe, M., De Oliveira, T., and Gascuel, O. (2018). Renewing felsenstein’s phylogenetic bootstrap in the era of big data. Nature, 556(7702):452–456.

Maddison, W. P. (1997). Gene Trees in Species Trees. Systematic Biology, 46(3):523–536. doi: 10.1093/sysbio/46.3.523.

Minh, B. Q., Schmidt, H. A., Chernomor, O., Schrempf, D., Woodhams, M. D., von Haeseler, A., and Lanfear, R. (2020). IQ-TREE 2: New Models and Efficient Methods for Phylogenetic Inference in the Genomic Era. Molecular Biology and Evolution, 37(5):1530–1534. doi: 10.1093/molbev/msaa015.

Moutsopoulos, I., Maischak, L., Lauzikaite, E., Vasquez Urbina, S. A., Williams, E. C., Drost, H.-G., and Mohorianu, I. I. (2021). noisyR: enhancing biological signal in sequencing datasets by characterizing random technical noise. Nucleic Acids Research, 49(14):e83–e83. doi: 10.1093/nar/gkab433.

Münkemüller, T., Lavergne, S., Bzeznik, B., Dray, S., Jombart, T., Schiffers, K., and Thuiller, W. (2012). How to measure and test phylogenetic signal. Methods in Ecology and Evolution, 3(4):743–756.

Piel, W. H., Chan, L., Dominus, M. J., Ruan, J., Vos, R. A., and Tannen, V. (2009). TreeBASE v. 2: A Database of Phylogenetic Knowledge. e-BioSphere 2009.

Price, M. N., Dehal, P. S., and Arkin, A. P. (2010). Fasttree 2 – approximately maximumlikelihood trees for large alignments. PLOS ONE, 5(3):1–10. doi: 10.1371/journal.pone.0009490.

Robinson, D. and Foulds, L. (1981). Comparison of phylogenetic trees. Mathematical Biosciences, 53(1):131–147. doi: 10.1016/0025-5564(81)90043-2.

Roch, S. (2006). A short proof that phylogenetic tree reconstruction by maximum likelihood is hard. IEEE/ACM Transactions on Computational Biology and Bioinformatics, 3(1):92–94. doi: 10.1109/TCBB.2006.4.

Rokas, A. and Carroll, S. B. (2006). Bushes in the tree of life. PLoS biology, 4(11):e352.

St. John, K. (2016). Review Paper: The Shape of Phylogenetic Treespace. Systematic Biology, 66(1):e83–e94. doi: 10.1093/sysbio/syw025.

Stamatakis, A. (2014). Raxml version 8: a tool for phylogenetic analysis and post-analysis of large phylogenies. Bioinformatics, 30(9):1312–1313.

Togkousidis, A., Kozlov, O. M., Haag, J., Höhler, D., and Stamatakis, A. (2023). Adaptive raxml-ng: Accelerating phylogenetic inference under maximum likelihood using dataset difficulty. Molecular Biology and Evolution, 40(10):msad227.

Townsend, J. P., Su, Z., and Tekle, Y. I. (2012). Phylogenetic Signal and Noise: Predicting the Power of a Data Set to Resolve Phylogeny. Systematic Biology, 61(5):835–835. doi: 10.1093/sysbio/sys036.

Trost, J., Haag, J., Höhler, D., Jacob, L., Stamatakis, A., and Boussau, B. (2024). Simulations of sequence evolution: how (un) realistic they are and why. Molecular biology and evolution, 41(1):msad277.

Vinh, L. S. and Von Haeseler, A. (2004). Iqpnni: moving fast through tree space and stopping in time. Molecular biology and evolution, 21(8):1565–1571.

Yang, Z. Molecular Evolution: A Statistical Approach. OUP Oxford, (2014). ISBN 9780191023309. URL https://books.google.gr/books?id=T-LoAwAAQBAJ.

