## Supplement for "Much Ado About Nothing: Accelerating Maximum Likelihood Phylogenetic Inference via Early Stopping to evade (Over-)optimization"

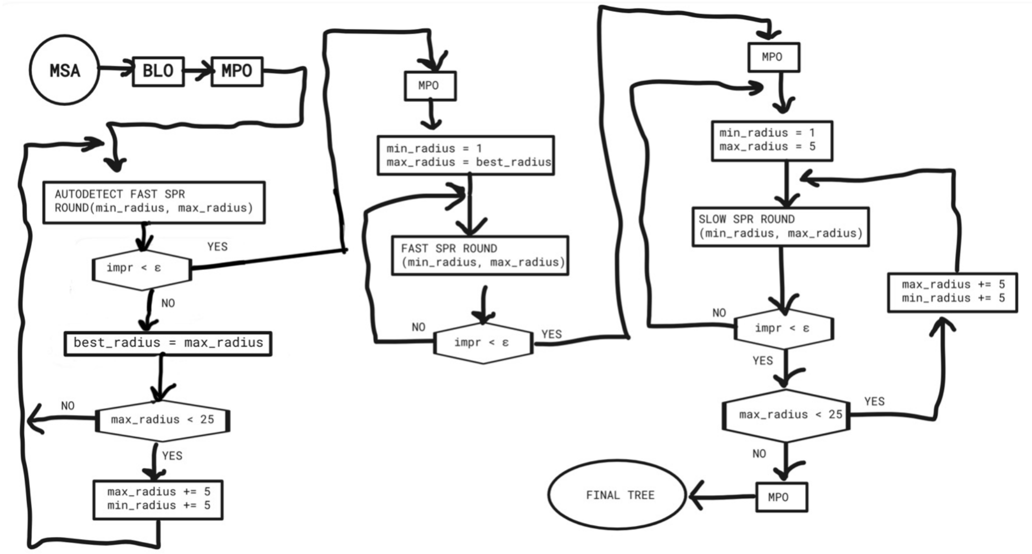


**Figure S1.** A schematic representation of the RAxML-NG v.1.2 search heuristic.

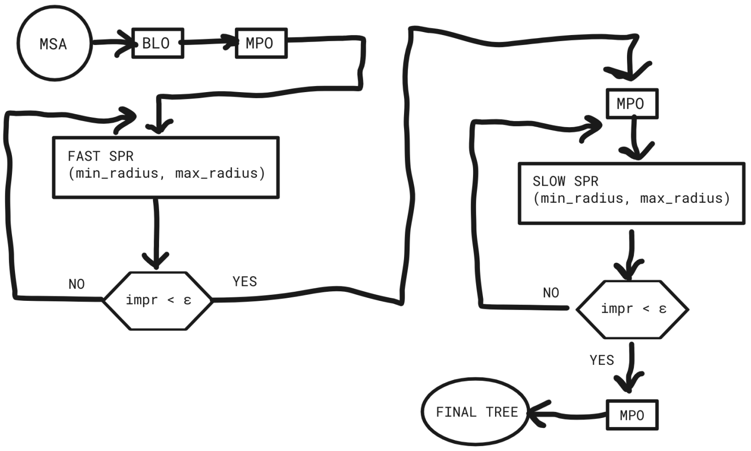


**Figure S2.** sRAxML-NG heuristic

### 1. Methods

In this section, we provide additional information on the implementation and specific considerations of the Early Stopping methods. This includes a detailed outline of the heuristic of the Simplified RAxML-NG version (sRAxML-NG), and specific boundary cases that required dedicated solutions in the KH-based versions.

#### 1.1. RAxML-NG v.1.2 vs sRAxML-NG

Figure S1 provides a schematic representation of the RAxML-NG v.1.2 tree search heuristic. This representation is simplified and omits unnecessary details, such as the exact implementation of Subtree Prune and Regraft (SPR) rounds as they are identical among the different versions. The purpose of this Figure is to provide an overview of the thoroughness of the original RAxML-NG heuristic and to highlight the limited as well as hard-to-interpret role of the $\epsilon$-thresholds in terminating the search. Additional technical details on the standard RAxML-NG heuristic can be found in Kozlov *et al.* (2018).

Branch-Length Optimization and Model-Parameter Optimization rounds are denoted by BLO and MPO, respectively. After each SPR round a full BLO round is conducted on the X best trees found during this round (see 2018 for details). For the sake of the simplicity we do not display this BLO step in Figure S1 but consider it as being an implicit part of the SPR round block in the diagram. The initial value of `best_radius` parameter is $5$.

The RAxML-NG v1.2 heuristic can be divided into three successive and different types of several SPR rounds, corresponding to the three closed loops of Figure S1. These SPR rounds are denoted by the following functions in the workflow:

1. `AUTODETECT FAST SPR ROUND(min_radius, max_radius)`: This function takes as input the minimum and maximum SPR radius parameters, which denote the minimum and maximum range (in terms of number of nodes away from the original branch where the subtree was pruned) of regrafting distances of an SPR move on the tree topology. These SPR rounds are essentially `FAST SPR` rounds (see below). The purpose of this initial sequence of SPR rounds is to automatically determine the `best_radius` parameter for the dataset at hand, which is used by `FAST SPR` rounds as the maximum regrafting distance. The algorithm successively increases the minimum and maximum radius parameters by $5$ units after each SPR round, and the sequence terminates when the log-likelihood improvement is less than the $\epsilon$-threshold or when the global maximum radius setting of $25$ units has been reached.

2. `FAST SPR ROUNDS(min_radius, max_radius)`: This function takes as input the minimum regrafting distance which is always set to 1 and the maximum regrafting distance as given by the `best_radius` parameter and determined via the preceding `AUTODETECT FAST SPR` rounds sequence. In `FAST SPR` rounds, RAxML-NG evaluates each tree topology generated by an SPR move (i.e., each subtree insertion) using the existing branch lengths. The sequence of consecutive `FAST SPR` rounds terminates when the log-likelihood improvement is less than the $\epsilon$-threshold.

3. `SLOW SPR ROUNDS(min_radius, max_radius)`: In analogy to the preceding SPR rounds, this function also takes as input the minimum and maximum regrafting distances, that are initially set to $1$ and $5$, respectively. The main difference between `FAST` and `SLOW SPR` rounds is that, during slow rounds, RAxML-NG evaluates each tree topology resulting from an SPR move by re-optimizing the lengths of the three adjacent branches around the insertion node. If, after an SPR round, the log-likelihood improvement is below the $\epsilon$-threshold, the minimum and maximum parameters are both increased by $5$; otherwise, the next SPR round the minimum and maximum radii remain $1$ and $5$, respectively. The sequence of `SLOW SPR` rounds is terminated when the log-likelihood improvement is below the $\epsilon$-threshold for five consecutive SLOW SPR round invocations, with increasing radius intervals (1-5, 6-10, 11-15, 16-20, 21-25) .

From the above description, it is evident that RAxML-NG v.1.2 can terminate when several distinct conditions are met. One critical factor is the $\epsilon$-threshold parameter. However, termination is also depends on other factors, such as the completion of three distinct sequences of increasingly thorough SPR rounds. Additionally, in the final sequence of SPR rounds, the algorithm terminates only if the log-likelihood improvement does not exceed the $\epsilon$-threshold for five consecutive SPR rounds with increasing minimum and maximum radii values. This thorough heuristic was designed to maximize the likelihood of the inferred topology to the best possible degree. As noted in the main text, Maximum Likelihood (ML) phylogenetic inference is an NP-hard optimization problem, as the number of possible topologies that need to be evaluated grows super-exponentially with the number of taxa in the input Multiple Sequence Alignment (MSA). Consequently, a superficial heuristic has a relatively high probability of becoming stuck in local optima.

In the main text, we argue that there are compelling reasons to not push the optimization to its limits. Therefore, we proposed four distinct stopping criteria that dynamically determine the $\epsilon$ parameter based on the noise present in the MSA and the convergence dynamics of the specific search process. The role of the $\epsilon$-threshold in terminating RAxML-NG v.1.2 is confounded with other stopping conditions. In addition, multiple unnecessary SPR rounds may be conducted despite the fact that the stopping criteria we propose may already indicate that execution should be halted. This emphasizes the need to simplify the RAxML-NG v.1.2 heuristic such that solely the $\epsilon$ parameter determines termination.

Figure S2 illustrates the sRAxML-NG heuristic. This heuristic only comprises two sequences of SPR rounds: one conducts `FAST SPR` and the other conducts `SLOW SPR` rounds. In both types of SPR rounds, the `min_radius` parameter is set to 1. The `max_radius` parameter has a fixed default value of 10, but users can modify this value using the `--spr-radius` option in the command line. Both sequences of SPR rounds terminate when the log-likelihood improvement after an SPR round is below the $\epsilon$-threshold. sRAxML-NG uses the same $\epsilon$-threshold as in RAxML-NG v.1.2, that is $10.0$ log-likelihood units. In the versions with adaptive stopping criteria, the $\epsilon$-threshold is either computed once at the beginning of the tree search (SN-based versions) or after each SPR round (KH-based versions).

##### 1.2. KH-based versions

As mentioned in the main text, certain boundary cases require special handling in the implementation of the KH-based versions. These boundary cases are listed below:

- If the log-likelihood improvement ($NL - L$) after an SPR round is negligible (down to numerical round-off error), it is expected that the standard deviation $\sigma_{KH}$ will also be negligible. Consequently, the quantity $\frac{NL-L}{\sigma_{KH}\sqrt{s}}$, used as an argument in the cumulative distribution function $\Phi(\cdot)$ of the standard normal distribution (see Sections 2.4 and 2.5 of the main text), becomes undefined, inducing numerical instability. To avoid this, when $NL - L < 10$, the algorithm exits the current sequence of SPR rounds. Essentially, the minimum value that the $\epsilon$-threshold can take in both KH-based methods is $10$. This prevents the KH-based methods from assigning a value to the $\epsilon$-threshold that is lower than the one used in RAxML-NG v.1.2, which has been empirically determined to suffice for infering high-quality trees.

- If the quantity $\frac{NL-L}{\sigma_{KH}\sqrt{s}}$ exceeds $3.5$, we assume that the Null Hypothesis is rejected (in both KH-based versions), as this value is typically the maximum in Z-score tables. In this case, we set $\epsilon := NL - L - 1$, and the algorithm proceeds to the next SPR round.
- Occasionally, $NL - L > 0$ may occur even if the topology remains unchanged before and after the SPR round, due to branch-length optimizations within and after the SPR round, again due to round-off error propagation. In such instances, we set $\epsilon := NL - L + 1$ and the algorithm exits the current SPR round sequence.

### 2. Commands

The stopping criteria are implemented in the `adaptive` branch[1] of the RAxML-NG repository. To disable the adaptive heuristic (Togkousidis et al., 2023) and run the standard version of RAxML-NG v.1.2, we use the `--adaptive off` command. sRAxML-NG can be invoked via `--extra simplified-on`. Stopping criteria are selected using the `--stopping-criterion` argument; available options are: `{sn-normal | sn-rell | KH | KH-mult}`, corresponding to the SN-Normal, SN-RELL, simple KH test, and KH test with multiple testing correction criteria. When stopping criteria are specified, the algorithm automatically executes the sRAxML-NG heuristic search, hence `--extra simplified-on` does not need to be specified explicitly. The commands we used to invoke each version and generate our results are the following (e.g., using $10$ parsimony starting trees):

**Standard RAxML-NG v.1.2:**

```
./raxml-ng-adaptive --adaptive off --threads 1 --msa {msa} --model {model} --tree pars{10}
--seed 0
```

**sRAxML-NG:**

```
./raxml-ng-adaptive --threads 1 --msa {msa} --model {model} --tree pars{10} --seed 0
--extra simplified-on
```

**SN-Normal version:**

```
./raxml-ng-adaptive --threads 1 --msa {msa} --model {model} --tree pars{10} --seed 0
--stopping-criterion sn-normal
```

**SN-RELL version:**

```
./raxml-ng-adaptive --threads 1 --msa {msa} --model {model} --tree pars{10} --seed 0
--stopping-criterion sn-rell
```

**KH version:**

```
./raxml-ng-adaptive --threads 1 --msa {msa} --model {model} --tree pars{10} --seed 0
--stopping-criterion KH
```

**KH multiple testing correction version:**

```
./raxml-ng-adaptive --threads 1 --msa {msa} --model {model} --tree pars{10} --seed 0
--stopping-criterion KH-mult
```

In the above commands, since the seed is fixed among all version invocations, and the modifications we introduced only affect the tree optimization stages, we are sure that all versions initiate the exact same trees.

### 3. Datasets

As mentioned in the main text, for our experiments we used $725$ empirical and $506$ simulated MSAs. Among the empirical datasets, $575$ are DNA and $150$ are amino acid (AA) MSAs. The empirical MSAs were sampled from the TreeBASE databse (Piel et al., 2009) and the simulated MSAs from the datasets used in a recent study on simulated data, conducted by our group and colleagues (Trost et al., 2024). For the analysis of the DNA MSAs we used the GTR+$\Gamma$ model (Tavaré, 1986), and for AA datasets we used the LG+$\Gamma$ model (Le and Gascuel, 2008). Information regarding the dimensions of the MSAs are provided in the scatter plots of Figure S3. The Pythia score distributions (Haag et al., 2022) of the sampled datasets are presented in Figure S4.

[1] https://github.com/togkousa/raxml-ng

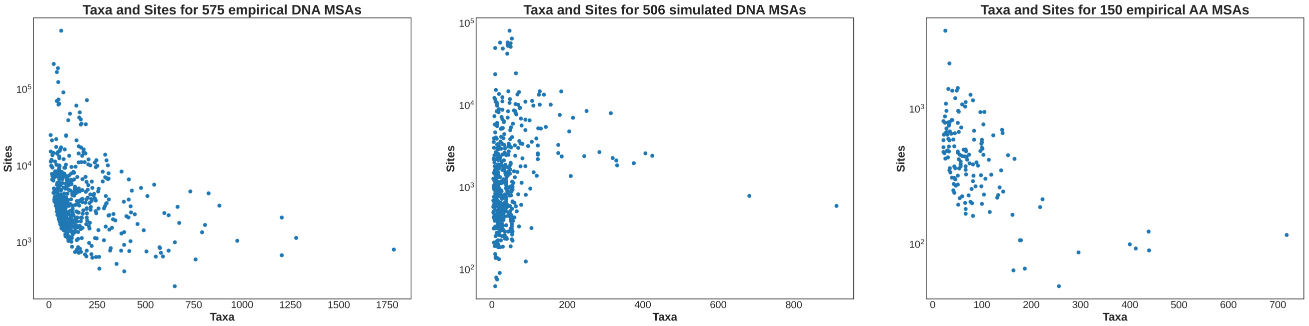


**(a)** **(b)** **(c)**

**Figure S3.** Scatter plot showing the number of taxa and sites for each dataset in **(a)** $575$ empirical DNA MSAs **(b)** $506$ simulated DNA MSAs and **(c)** $150$ empirical AA MSAs.

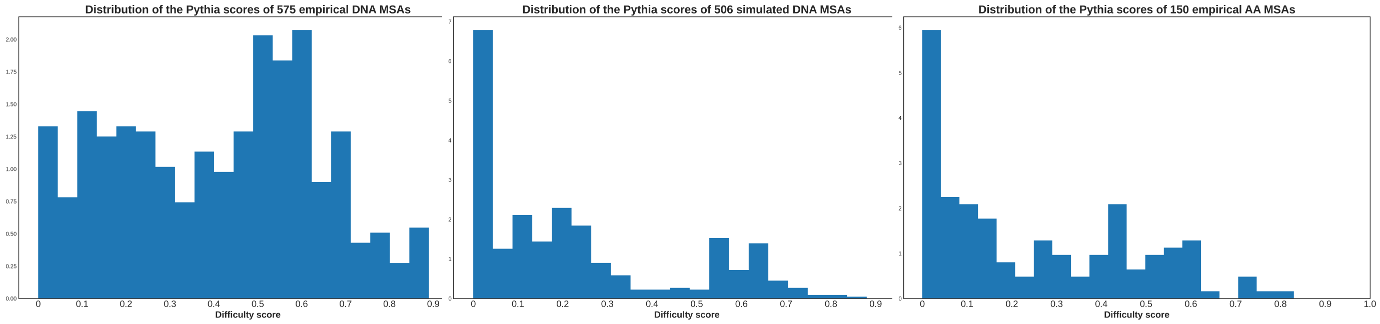


**(a)** **(b)** **(c)**

**Figure S4.** Distributions of Pythia scores of **(a)** $575$ empirical DNA, **(b)** $506$ simulated DNA, and **(c)** $150$ empirical AA MSAs.

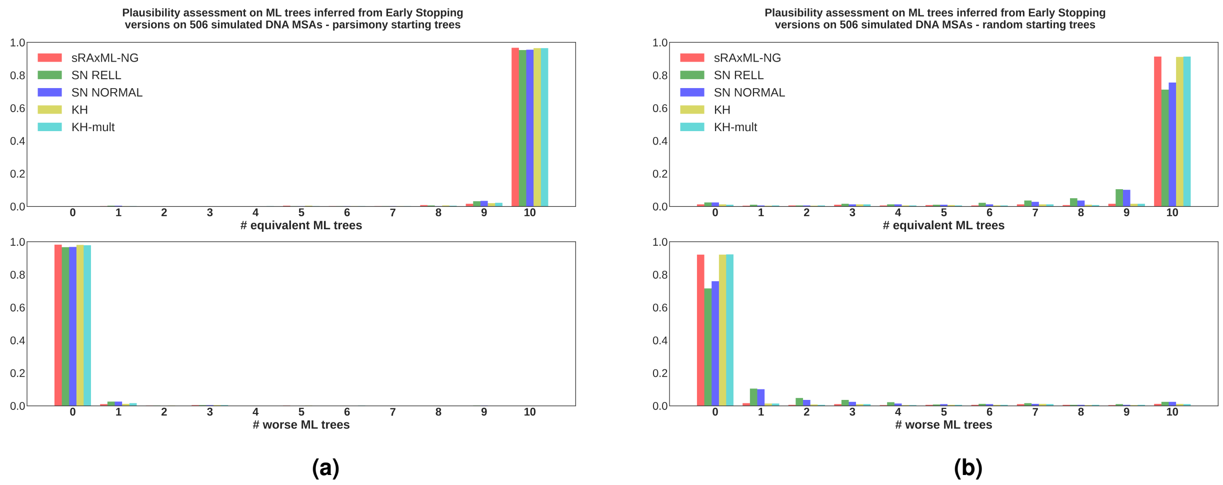


**Figure S5. (a)** Results from the plausibility tests conducted on the ML trees inferred from Early stopping criteria on $506$ simulated DNA MSAs. For each dataset, all RAxML-NG versions conduct $10$ independent tree inferences using (the same) set of parsimony starting trees. The ML trees inferred using Early Stopping criteria are compared to the optimal ML tree inferred from RAxML-NG v.1.2 using the KH test in a pair-wise manner. The horizontal axes shows the number (out of $10$) of the ML trees inferred by each version that are statistically equivalent (top) and worse (bottom) than the best-known ML tree inferred via RAxML-NG v.1.2. The vertical axes shows the fraction of MSAs whose analysis yielded the corresponding number of equivalent or worse ML trees, respectively. **(b)** Identical experimental setup as under (a), with the sole difference being that we use $10$ random starting trees instead of $10$ parsimony starting tree.

### 4. Supplementary Results

In the main text, we only presented results for empirical and simulated DNA datasets. Here, we provide the results from analyzing the AA datasets, as well as additional plots from the analysis of the DNA MSAs which were omitted in the main text.

#### 4.1. Results on DNA datasets

Figure S5 illustrates the distribution of the number of statistically equivalent and worse ML trees inferred from Early Stopping versions across simulated DNA datasets. Instances where Early Stopping versions infer statistically better ML trees are discussed below. This plot is analogous to Figure 1 in the main text but refers to simulated MSAs instead of empirical ones. As mentioned in the main text, simulated MSAs are generally easier to analyze. This explains why for all Early Stopping versions, the majority of the inferred ML trees are plausible. Additionally, we observe a decrease in the performance of the SN-based versions when using random starting trees, although this decrease is not as pronounced as for empirical MSAs.

Regarding the results from parsimony starting trees (Figure S5a), all Early stopping versions infer at least one plausible tree on all $506$ simulated MSAs ($100\%$ of the cases). Additionally, the number of MSAs where the SN-based methods infer $10$ plausible trees is $483$ for the SN-Normal and $482$ for the SN-RELL version ($99\%$ of the cases). The corresponding numbers for the remaining methods are $489$ for sRAxML-NG and $488$ for the KH-based versions ($96\%$). Concerning the rare instances where the Early Stopping methods infer statistically better trees than RAxML-NG v.1.2., sRAxML-NG and the KH-based versions infer at least one better tree on $9$ MSAs, and the SN-based versions on $8$ datasets.

Regarding the results on random starting trees (Figure S5b), in $98\%$ of the cases all Early stopping versions infer at least one plausible tree. sRAxML-NG and the KH-based versions infer $10$ plausible trees on $462$ simulated MSAs ($91\%$), while the corresponding numbers for SN-Normal are $382$ ($75\%$) and for SN-RELL are $360$ ($71\%$). Further, the KH-based versions infer at least one statistically better tree on $6$ MSAs, the sRAxML-NG on $5$, the SN-Normal on $4$ and the SN-RELL on $3$ MSAs.

Figure S6 illustrates the speedups attained from early stopping on both, empirical, and simulated DNA MSAs, when random starting trees are used. Figures S6a and S6b are analogous to Figures 3 and 4 of the main text, with no substantial differences. Figure S6a illustrates the distributions of runtime improvements when Early Stopping versions are compared to RAxML-NG v.1.2, while Figure S6b highlights the additional speedup attained via the SN-based and KH-based versions compared to sRAxML-NG.

Figure S7 illustrates the distributions of Robinson-Foulds (RF) distances between the best ML trees inferred from each RAxML-NG version and the reference tree topologies when random starting trees are used. For simulated

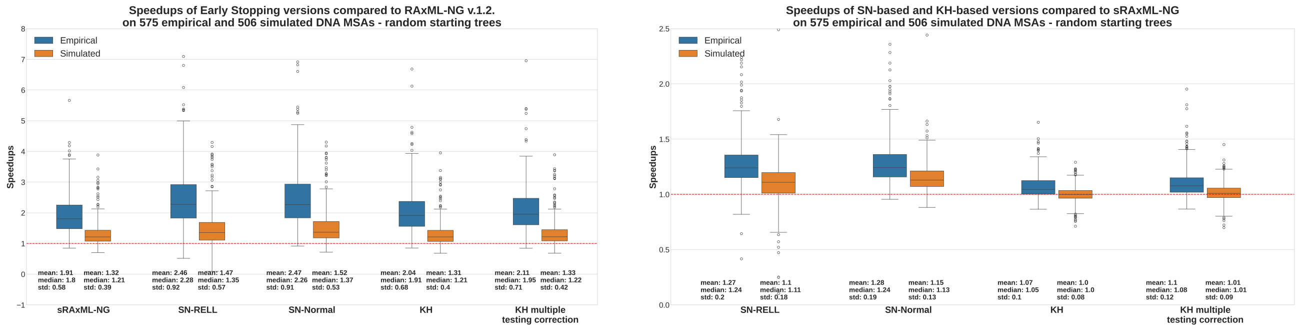


**(a)** **(b)**

**Figure S6.** **(a)** Speedup distributions of Early Stopping versions relative to RAxML v.1.2 on $575$ empirical and $506$ simulated DNA MSAs. The speedups refer to sequential runtimes using random starting trees. The dashed line at the bottom corresponds to a speedup of $1$x. **(b)** Speedup distributions of SN-based and KH-based RAxML-NG versions relative to sRAxML-NG on $575$ empirical and $506$ simulated DNA MSAs. The speedups refer to sequential runtimes using random starting trees. The dashed line at the bottom corresponds to a speedup of $1$x.

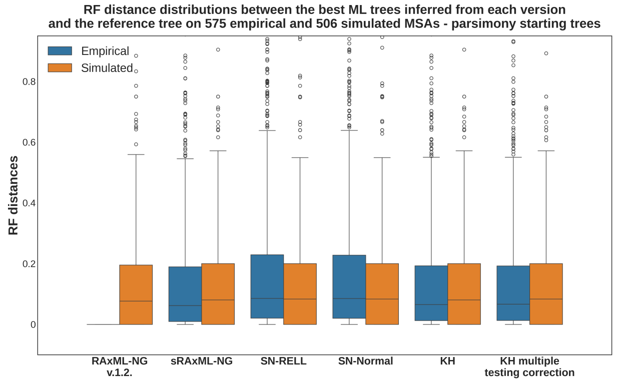


**Figure S7.** Distributions of relative RF distances between the best ML trees inferred from each RAxML-NG version and the corresponding reference tree topologies, where each version conducted $10$ independent tree inferences on $575$ empirical and $506$ simulated DNA MSAs using random starting trees. On simulated MSAs, the reference tree topology is the "true" tree used to generate the sequences. For empirical MSAs, the reference topology is the best ML tree inferred by RAxML-NG v.1.2. This also explains the absence of a distribution for empirical MSAs in the RAxML-NG v.1.2 column.

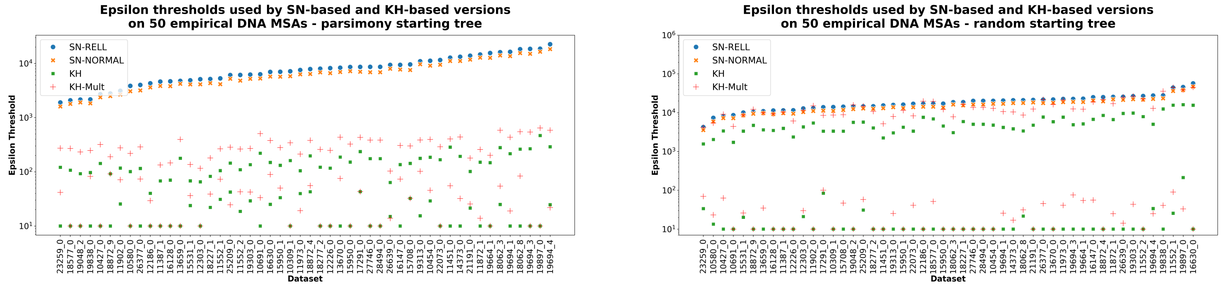


**(a)** **(b)**

**Figure S8.** Convergence $\epsilon$-thresholds used by Early Stopping versions for a single tree inference on $50$ empirical DNA MSAs, using a parsimony starting tree. The datasets are sorted based on the $\epsilon$-thresholds determined in the SN-RELL version. For the KH-based methods, we only plot the maximum and minimum $\epsilon$-thresholds used throughout the tree inference. **(b)** Same description as in (a), however each version uses a single random starting tree instead of a parsimony starting tree. The datasets on the x-axis are the same in Figures (a) and (b).

MSAs, we observe no substantial differences between the relative RF distance distributions in the current figure and those in Figure 2 of the main text, which depicts analogous results, but for parsimony starting trees. As already mentioned in the main text, all relative RF distance distributions between the best ML trees inferred by all versions and the true tree on simulated MSAs (when using both, random, and parsimony starting trees), have the same average of $0.13$. On the other hand, the RF distributions for empirical MSAs show higher RF distances compared to those in Figure 2 of the main text. As noted in the main text, drawing definitive conclusions about the RF distance distributions of empirical MSAs is challenging because the true tree topology is unknown. We compare the topologies inferred by the Early Stopping versions against the topology inferred by RAxML-NG v.1.2. The slight shift to higher RF distances when random starting trees are being used is rather expected, since the starting points for tree inferences are evidently worse.

Finally, in Figure S8, we present the $\epsilon$-threshold values used by each Early Stopping version for a single tree inference on a randomly selected subset of $50$ (out of $575$) empirical DNA MSAs. Figure S8a shows the $\epsilon$-thresholds when using a parsimony starting tree, while Figure S8b when using a random starting tree. This plot illustrates the range of $\epsilon$ values used by each Early Stopping version. For the KH-based methods, where the $\epsilon$ value is recomputed after each SPR round and adjusted based on the respective convergence dynamics and noise in the per-site log-likelihood values, only the minimum and maximum values are shown.

As already reported in the main text, the SN-based stopping criteria yield larger thresholds, ranging from hundreds to thousands of log-likelihood units, depending on how well the starting tree on which they are computed fits the input data. If we compare the two plots, we observe that the $\epsilon$-thresholds of SN-based methods are substantially larger on a random starting trees. Despite these large convergence thresholds, SN-based methods typically find statistically plausible trees in the majority of cases, especially when parsimony starting trees are used. This is because the vast tree space, which grows super-exponentially with the number of taxa, contains multiple search paths that hill-climbing heuristics can follow to reach local optima. SPR moves, being highly effective topological moves, often result in substantial improvements in the log-likelihood score. The worse the initial log-likelihood score before the SPR round, the greater the subsequent improvement after the SPR round. Consequently, the large $\epsilon$-thresholds set by SN-based methods are frequently exceeded by these improvements (especially by the improvement of the first SPR round). When starting from a parsimony tree, one or two SPR rounds are often sufficient for Early Stopping versions to infer trees that are statistically equivalent to the best ML tree inferred by RAxML-NG v1.2.

For the KH-based versions, the KH method with multiple testing correction yields higher $\epsilon$-thresholds than the simple KH method, especially during the initial SPR rounds, where hundreds of SPR topologies $(N_t)$ improve the log-likelihood score of the input tree (see eq. (3) in the main text). This illustrates the advantage of a dynamic approach that adapts the $\epsilon$ value as a function of the data and intermediate trees. Moreover, we observe that the minimum $\epsilon$ value for both versions is $10$. This constraint arises from the necessary numerical restrictions in the implementation of the KH test (see Section 1.2) to ensure that the $\epsilon$ value computed by the KH-based methods is not lower than the threshold used in RAxML-NG v1.2, which is $10$.

##### 4.2. Results on AA datasets

Figure S9 illustrates the distribution of the number of statistically equivalent and worse ML trees inferred from Early Stopping versions across $150$ AA datasets. We do not detect instances where Early Stopping versions infer statistically better ML trees than RAxML-NG v.1.2 on AA datasets. This figure is analogous to Figures S5 for simulated

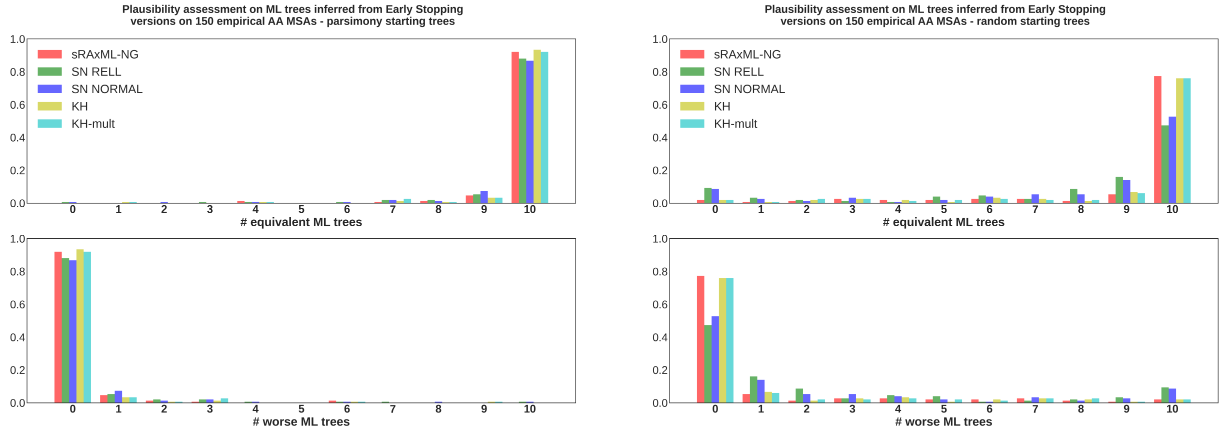


**(a)** **(b)**

**Figure S9. (a)** Results from the plausibility tests conducted on the ML trees inferred from Early stopping criteria on $150$ empirical AA MSAs. For each dataset, all RAxML-NG versions conduct $10$ independent tree inferences using (the same) set of parsimony starting trees. The ML trees inferred using Early Stopping criteria are compared to the optimal ML tree inferred from RAxML-NG v.1.2 using the KH test in a pair-wise manner. The horizontal axes shows the number (out of $10$) of the ML trees inferred by each version that are statistically equivalent (top) and worse (bottom) than the best-known ML tree inferred via RAxML-NG v.1.2. The vertical axes shows the fraction of MSAs whose analysis yielded the corresponding number of equivalent or worse ML trees, respectively. **(b)** Identical experimental setup as under (a), with the sole difference being that we use $10$ random starting trees instead of $10$ parsimony starting tree.

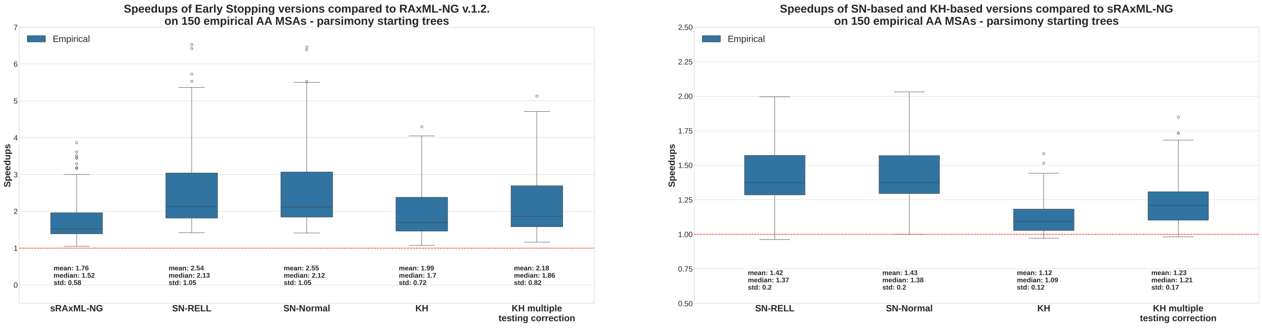


**(a)** **(b)**

**Figure S10.** **(a)** Speedup distributions of Early Stopping versions relative to RAxML v.1.2 on $150$ empirical AA MSAs. The speedups refer to sequential runtimes using parsimony starting trees. The dashed line at the bottom corresponds to a speedup of $1$x. **(b)** Speedup distributions of SN-based and KH-based RAxML-NG versions relative to sRAxML-NG on $150$ empirical AA MSAs. The speedups refer to sequential runtimes using parsimony starting trees. The dashed line at the bottom corresponds to a speedup of $1$x.

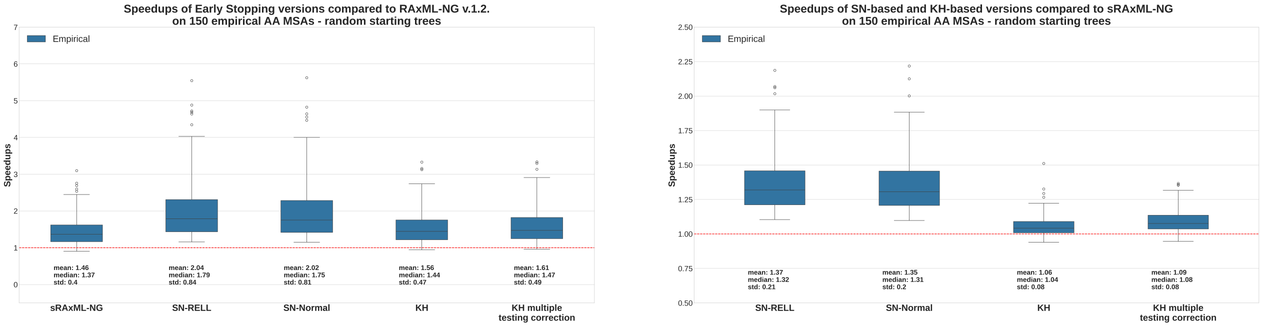


**(a)** **(b)**

**Figure S11.** **(a)** Speedup distributions of Early Stopping versions relative to RAxML v.1.2 on $150$ empirical AA MSAs. The speedups refer to sequential runtimes using random starting trees. The dashed line at the bottom corresponds to a speedup of $1$x. **(b)** Speedup distributions of SN-based and KH-based RAxML-NG versions relative to sRAxML-NG on $150$ empirical AA MSAs. The speedups refer to sequential runtimes using random starting trees. The dashed line at the bottom corresponds to a speedup of $1$x.

DNA and to Figure 1 (in the main text) for empirical DNA datasets. The results are analogous to those obtained for empirical DNA MSAs.

Regarding the results from parsimony starting trees (Figure S9a), sRAxML-NG and the KH-based versions infer at least one plausible tree on all $150$ AA MSAs ($100\%$ of the cases), while the SN-based methods do so on $149$ MSAs. The number of MSAs in which sRAxML-NG infers $10$ plausible trees is $138$ ($92\%$); the corresponding number for SN-based versions is $130$, for SN-Normal ($87\%$), and $132$ for SN-RELL version ($88\%$). For KH-based versions the numbers are $140$ ($93\%$) for the simple KH test and $138$ ($92\%$) AA MSAs for the KH test with multiple testing correction.

As expected, the results from random starting trees (Figure S9b) show a performance decrease for SN-based versions. In $98\%$ of the cases, sRAxML-NG and the KH-based versions infer at least one plausible tree, while the corresponding rate for the SN-based versions is $91\%$. Further, sRAxML-NG infers $10$ plausible trees on $116$ AA MSAs ($77\%$), and the KH-based versions on $114$ ($76\%$). The SN-Normal infers $10$ plausible trees on $79$ ($53\%$), and the SN-RELL on $71$ ($47\%$) AA datasets.

Figures S10 and S11 illustrate the distributions of runtime improvements on empirical AA MSAs. Figures S10a and S10b refer to speedups of Early Stopping versions compared to RAxML-NG v.1.2, while Figures S10b and S11b refer to speedups of SN-based and KH-based versions compared to sRAxML-NG. The speedup distributions are analogous to those on empirical DNA MSAs, indicating (in combination with Figure S9) that our stopping criteria can also be applied to AA datasets.

Finally, Figure S12 shows the distributions of relative RF distances between the best ML trees inferred from each Early Stopping version and RAxML-NG v.1.2, when analyzing the $150$ AA datasets.

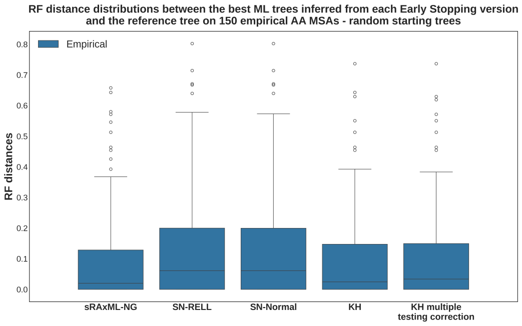


**(a)**

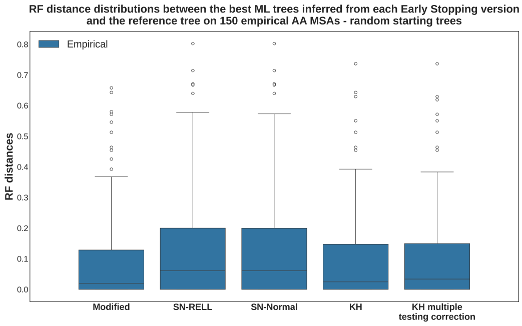


**(b)**

**Figure S12. (a)** Distributions of relative RF distances between the optimal ML trees inferred from each Early Stooping version and the optimal ML tree inferred from RAxML-NG v.1.2, on $150$ empirical AA datasets. Each version conducted $10$ independent tree inferences on the $150$ AA MSAs using the same set of parsimony starting trees. **(b)** Same description as in (a), but $10$ random starting trees are used instead.

#### Acknowledgment

This study was financially supported by the Klaus Tschira Foundation and by the European Union (EU) under Grant Agreement No 101087081 (Comp-Biodiv-GR).

Funded by
the European Union

#### References

Haag, J., Höhler, D., Bettisworth, B., and Stamatakis, A. (2022). From easy to hopeless—predicting the difficulty of phylogenetic analyses. *Molecular Biology and Evolution*, 39(12):msac254.

Kozlov, A. *Models, Optimizations, and Tools for Large-Scale Phylogenetic Inference, Handling Sequence Uncertainty, and Taxonomic Validation.* PhD thesis, Karlsruhe Institute of Technology, (2018).

Le, S. Q. and Gascuel, O. (2008). An improved general amino acid replacement matrix. *Molecular biology and evolution*, 25(7):1307–1320.

Piel, W. H., Chan, L., Dominus, M. J., Ruan, J., Vos, R. A., and Tannen, V. (2009). TreeBASE v. 2: A Database of Phylogenetic Knowledge. *e-BioSphere 2009*.

Tavaré, S. (1986). Some probabilistic and statistical problems in the analysis of dna sequences. *Lect Math Life Sci (Am Math Soc)*, 17:57–86.

Togkousidis, A., Kozlov, O. M., Haag, J., Höhler, D., and Stamatakis, A. (2023). Adaptive raxml-ng: Accelerating phylogenetic inference under maximum likelihood using dataset difficulty. *Molecular Biology and Evolution*, 40(10):msad227.

Trost, J., Haag, J., Höhler, D., Jacob, L., Stamatakis, A., and Boussau, B. (2024). Simulations of sequence evolution: how (un) realistic they are and why. *Molecular biology and evolution*, 41(1): msad277.
